# Evolution and phylogenetic distribution of *endo*-α-mannosidase

**DOI:** 10.1101/2022.12.21.521504

**Authors:** Łukasz F. Sobala

## Abstract

While glycans underlie many biological processes, such as protein folding, cell adhesion and cell-cell recognition, deep evolution of glycosylation machinery remains an understudied topic. N-linked glycosylation is a conserved process in which mannosidases are key trimming enzymes. One of them is the glycoprotein *endo*-α-1,2-mannosidase which participates in the initial trimming of mannose moieties from an N-linked glycan inside the *cis*-Golgi. It is unique as the only endo-acting mannosidase found in this organelle. Relatively little is known about its origins and evolutionary history; so far it was thought to occur only in vertebrates. Here I perform a taxon-rich bioinformatic survey to unravel the evolutionary history of this enzyme, including all major eukaryotic clades and a wide representation of animals. I found the endomannosidase to be vastly more widely distributed in animals than previously thought and in fact present in almost all eukaryotic clades. I tracked protein motif changes in context of the canonical animal enzyme. Additionally, my data show that the two canonical versions of endomannosidase in vertebrates, MANEA and MANEAL, arose at the second round of the two vertebrate genome duplications and indicate presence of a third protein, named here CMANEAL. Finally, I describe a framework where N-glycosylation co-evolved with complex multicellularity. A better understanding of the evolution of core glycosylation pathways is pivotal to understanding biology of eukaryotes in general, and the Golgi apparatus in particular. This systematic analysis of the endomannosidase evolution is one step towards this goal.

## Introduction

Glycans are integral to the structures of most biomolecules and participate in many processes that are instrumental to life. The enormous variety of the possible monomers, compounded by the large number of ways they may be covalently linked, made them uniquely suited for what multicellular organisms and cells within them have to do: differentiate self from non-self^1^. The best known examples of the effects of glycan differences are the human blood group system, ABO^2^, and the 3α-galactobiose epitope – the major reason for the rejection of pig-to-human xenotransplants^3^. Sialic acids, which are usually attached at the end of a glycan chain, often mediate host-parasite interactions^4^, such as immunity evasion^5^ or viral entry^6^. Parasitic helminths express glycans which co-evolved with host glycans^7^ to help evade the immune response. Tumors are characterized by altered glycosylation patterns^8^. Cellular adhesion is also affected by glycosylation – in this process, the physical properties of glycans and glycoproteins play a role. It is thought that in the last common ancestor of animals^9,10^, dystroglycan was present and functional in cell-ECM (extracellular matrix) adhesion.

In multicellular organisms, cells are under constant pressure to be consistent in their proteoform output. If they are found by other cells as non-self, they risk being being eliminated, akin to xenotransplants. This and other factors led animals to evolve exquisite ways of chaperoning protein folding and quality control. Protein N-glycosylation in the endoplasmic reticulum is one of the chaperoning pathways and depends on concerted activity of many biosynthetic enzymes, lectins and transporters. Oligosaccharyltransferase (OST)-based N-glycosylation is present in many eukaryotic species and some archaea^11^. The evolution of the OST complex central to this process has been studied in some detail already and detecting it in archaea was a finding which strongly supports the Asgard archaea clade as the closest relatives of all eukaryotes^12^.

Among many transporters and enzymes responsible for glycome formation, the function of the Golgi endo-α-1,2-mannosidase is to enable a higher proportion on N-glycans to be processed to complex structures, adding “polish” to the N-glycans. It is localized to *cis*-Golgi and the ER-Golgi intermediate compartment (ERGIC)^13^. As the only endo-acting mannosidase in this organelle, it is an enzyme with a unique catalytic activity. In this work I focus on its pedigree and evolution of its function in the context of changing life histories. I will make an attempt at answering the question of why its activity might be necessary for cells.

The endomannosidase is classified by the Carbohydrate Active Enzyme (CAZy) database (http://www.cazy.org/) in family GH99^14^. Its discovery^15^ explained previous observations of biosynthetic N-glycan trimming despite inhibition of glucosidase II^16,17^ and Golgi mannosidase I^18,19^. *In vivo*, the animal endomannosidase catalyzes the removal of di-, tri- or tetrasaccharide from the long arm of the precursor N-glycan attached to a glycoprotein (Fig. 1). The enzyme has the highest affinity towards a 3αGlc-2αMan-2αMan-Man minimal substrate, releasing 3αGlc-Man and 2α-mannobiose as the products (Fig. 1). The affinity of the Golgi endomannosidase to di- and triglucosylated substrates, as well as mannosylated substrates, is lower. Indeed, the bovine endomannosidase was found to restrict its processing to monoglucosylated substrates^20^. The reaction mechanism of GH99 was recently described in detail^21^ and the catalytic residues in the human MANEA (gene *MANEA* named after: MANnosidase Endo-Alpha) enzyme are thought to be E404 and E407 – identical to bacterial GH99 enzymes.

**Fig. 1:**
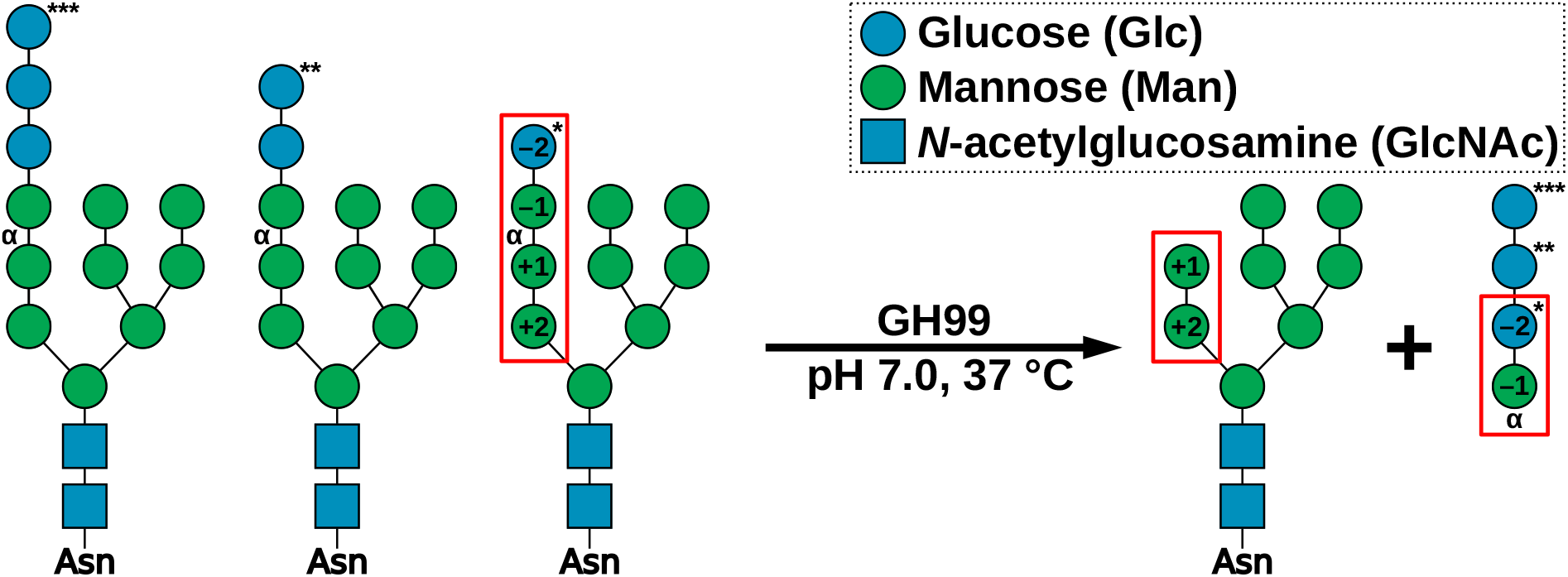
Reaction catalyzed by the Golgi endomannosidase. Corresponding substrate and product variants are marked with the same number of asterisks. The minimal substrate and minimal products are in red rectangles. Subsite positions^22^ are written inside the symbols.

According to sequence similarity, the GH99 glycoside hydrolase family is most closely related to GH71^23^. Proteins from this family are present in many *Actinobacteria*^24^ and fungi and are involved in glucan degradation^25^. Their similarities do not end with sequences, as GH71 enzymes (mutanases) share with GH99 a tetrasaccharide minimal substrate and *endo*activity^26^. Unlike GH99, the reaction mechanism of GH71 domain proteins was not studied and the catalytic residues are unknown. The second most closely related family is GH25^23^, which includes the ubiquitous lysozyme, also present in bacteria^27,28^.

Despite the importance of glycosylation, the evolutionary trajectories which shaped this pathway were rarely studied in detail, although notable comparative studies exist^29–32^. In particular, the evolutionary history of the endomannosidase remains obscure: so far, only two works were published. In an early taxonomic survey of biochemical activity^33^, Dairaku and Spiro concluded that the endomannosidase was a relatively recent addition to the N-glycan trimming enzyme repertoire, absent from plants, *Tetrahymena, Trypanosoma, Leishmania* and yeast. All vertebrates were found to possess endomannosidase activity. The only invertebrates investigated in this study were arthropods and molluscs, and it was stated that non-chordates lack this activity with a “conspicuous” exception of molluscs. Sobala^34^ conducted a limited phylogenetic study where it was stated that while many insects and other invertebrates do contain the endomannosidase, sponges and ctenophores lack this protein. It was also argued that bony vertebrates (Euteleostomi) contain two separate genes encoding for GH99 domain proteins, *MANEA* and *MANEAL*, while cartilaginous fishes (Chondrichthyes) possess only one gene. However, the taxon sampling in those two studies were limited. The functions of GH99 domain proteins from less closely related organisms was never studied. Possible functional variation of various clusters of GH99 domain proteins was mentioned in a recent review^35^, but all eukaryotic proteins formed a single cluster and the evolution of their sequence motifs was not investigated further.

The availability of sequencing data I have today allows me to re-analyze and extend these findings under a greater evolutionary context and a wider taxon sampling. Here I perform a taxon-rich bioinformatic survey to decipher the origin and the evolutionary history of GH99 domain proteins. I found that contrary to previous suggestions, the endomannosidase is widely distributed in eukaryotes, and in animals it is not limited to vertebrates but almost ubiquitous. Phylogenetic analyses enabled me to track the duplication history of the endomannosidase and discover the existence of a previously unrecognized vertebrate ortholog, present only in certain fish. In this article I will use the amino acid numbering of the human MANEA protein (UniProt ID **Q5SRI9**) as a reference when discussing motif evolution and protein variants from other species.

## Results

### Endo-α-1,2-mannanase/mannosidase is widespread in eukaryotes and has two main specificities

I found GH99 domain proteins in many clades of bacteria, archaea and eukaryotes. In the LukProt database (see Materials and Methods), the only eukaryotic clades that did not appear to have any proteins with the GH99 domain were Colponemidae, Peronosporomycetes and Apicomplexa, all within the broad Diaphoretickes group (Fig. 2). Due to data availability, my taxon sampling was very low for Colponemidae, which makes this conclusion tentative. In Podiata^36^, a clade that contains animals, GH99 is present in some form in all except Breviatea, Ichthyosporea, Acanthoecida and ctenophores. The low number of sampled breviates (3) leaves open the possibility that they possess proteins with this domain, but the other clades likely suffered secondary GH99 losses. These secondary losses are common in eukaryotes: for example, only two species of Streptophyta have the endomannosidase and none of them are land plants. However, a GH99 protein was found in *Cylindrocystis brebissonii* from a green algae clade Zygnematophyceae, which is sister to land plants.

**Fig. 2:**
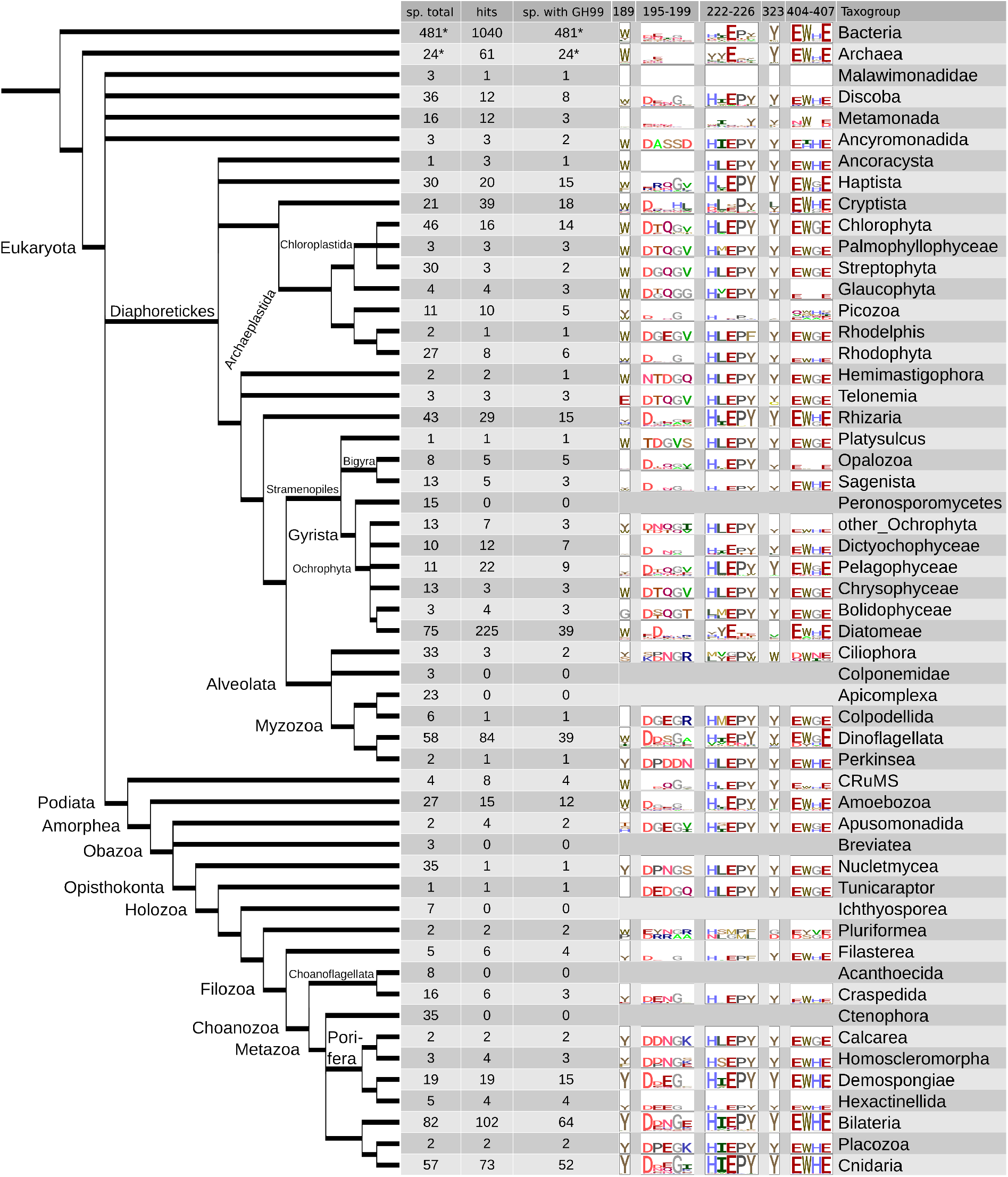
Analysis of the prevalence of GH99 domain proteins in the LukProt+Picozoa+*Txikispora* database and the evolution of its major sequence motifs. Asterisks denote taxogroups^37^ where the absolute prevalence could not have been estimated because of the unknown number of searched species (sp.). Sequence logos created using ggseqlogo^38^. Only sequences remaining after multiple rounds of phylogeny-based decontamination were used to build the sequence logos. In the table, the proportion of species with the GH99 domain is slightly lower than in the full database because the clustering method used (cd-hit at 95% identity) eliminated some highly similar species.

The GH99 protein family underwent a considerable expansion and diversification in diatoms (Diatomeae), some of which have a high number of hits per species. An extreme example is *Nitzschia sp*. RCC80^39^, in which 21 predicted proteins (clustered at 95% identity) were found. GH99 protein sequences from diatoms form two well-supported major clades (Supplementary Fig. S1).

I assumed that the predicted specificity of the endomannosidase may be used as a proxy pointing to its usage by cells. In most clades the endomannosidase is specific towards a Man4 tetrasaccharide minimal substrate. The preference is determined by residue 189, in most organisms a tryptophan^40^ (Fig. 3) which interacts hydrophobically with the −2 mannose (Fig. 3B). A tyrosine at position 189 switches the −2 sugar residue preference to Glc by forming a water-mediated hydrogen bond with the −2 sugar hydroxyl group^35,41^ (Fig. 3A).

**Fig. 3:**
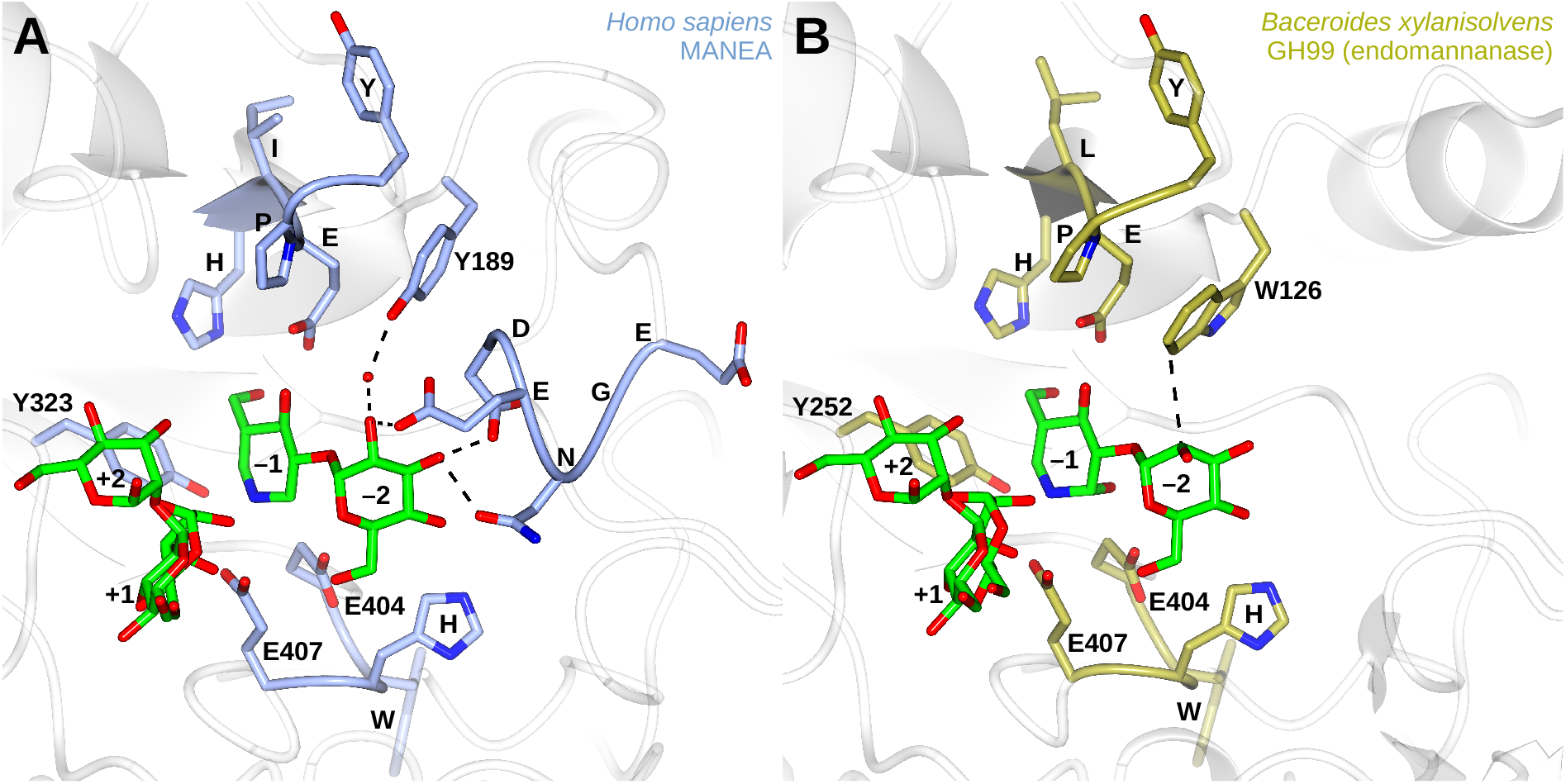
Comparison of the main motifs of (A) human GH99 (PDB ID: **6ZFA**) and bacterial GH99 (PDB ID: **5M03**) and its mode of binding of the substrate. All five investigated motifs can be seen in (A). Sugars are numbered according to the subsite^22^ they occupy. Figure prepared in ccp4mg^42^.

Among the GH99 sequences found in bacteria, only 19% contained Y189 and 63% had W189. The proportion in Archaea was even lower: 3% vs 85%. It is known that Bacteroidetes use the endomannanase to digest yeast mannan^43^. The endomannanase activity, however, predates divergence of fungi over 1 billion years ago^44^. I posit that the endomannosidase/endomannanase activity evolved much earlier and organisms first utilized this activity to remove recalcitrant branches of various exogenous mannose-containing glycans. This would make the smaller oligosaccharides available for downstream exoglycosidases which would then release monosaccharides ready for use in the cell’s own metabolism. It is likely that first eukaryotes used their GH99 endomannosidase/endomannanase in a similar way to bacteria and it became a part of the Golgi N-glycan processing pathway only in some lineages. Partial, reconstructed GH99 proteins from Ancyromonadida, thought to be a very early diverging lineage of eukaryotes^45^, contain all the crucial active site residues, as well as W189, signifying the ancient origin of this catalytic activity.

In Amorphea, within which animals are contained and whose sister group is CruMs^45^ (Fig. 2), I observed a switch to Y189. In none of the GH99 sequences from CRuMS (4 taxa sampled) Y189 was present, but a significant proportion (4 sequences, 27%) of hits from Amoebozoa have this feature (Supplementary File S11). Within Obazoa, a sister clade to Amoebozoa, only Opisthokonta retained Y189 and among the opisthokonts, the only nucletmycean that contains it is the recently sequenced *Parvularia atlantis*^46^. No true fungi possess the GH99 domain. Among the closest relatives of animals, the facultatively multicellular holozoans, Tunicaraptor and Pluriformea have various residues at site 189, but Filasterea seems to be the first clade where Y189 is prevailing (4 sequences, 67%). In craspedid choanoflagellates this Y/W proportion is 83% vs 0% and in non-ctenophore animals it is 77% vs 0%. Thus, it appears that Y189 is a feature of the last common ancestor of Choanozoa.

The W189Y switch also happened in Rhizaria (Fig. 2) in the Chlorarachniophyta group. For example, the model rhizarian *Bigelowiella natans* as well as *Chlorarachnion reptans* have two GH99 sequences, one with W189 and one with Y189. These sequences have different 195-199 loop structures, further pointing to differences in their substrate specificity. In these species, W189 is coupled with an **E**WG**E** active site motif and Y189 with **E**WH**E**. The W189/**E**WG**E** sequences are most similar to GH99 sequences from Cryptista and Chlorophyta, suggesting they are descendants of the sequence from the phototrophic endosymbiont^47^ (Supplementary Figure S1). The Y189/**E**WH**E** sequences come from the rhizarian host. Another group where the switch also happened may be Picozoa, although sequences from them might be less reliable as sequences from these species come mostly from single-cell genomes.

The 195-199 motif **D**[E/D]N**G**E seems to be a common feature in eukaryotic GH99 proteins, with D196 and G198 conserved in many clades. The notable exceptions are Haptista where only G198 is conserved, Cryptista with a partial consensus motif, and Diatomeae with only D196 conserved. The 222-226 motif H[I/L]**E**PY is retained in all clades except Diatomeae and Metamonada. Y323, which was suggested to be a potential nucleophile residue^48^, is indeed quite conserved but does not seem necessary for catalysis: for instance it was substituted for F323 in full length MANEAL proteins in certain fish species. Finally, the consensus catalytic site motif is **E**W[H/G]**E** across all domains of life (except Picozoa, for which the source sequences might be less accurate).

Building upon the understanding of the structure and motif evolution in GH99, I compared this domain to its closest CAZy family, GH71, also present in bacteria and eukaryotes. The high overall prevalence of GH71 and GH99 suggests they diverged before Archaea separated from bacteria. I include a manually constructed alignment of human GH99 and a GH71 based on an AlphaFold structure^49^ of *Emericella nidulans* mutanase (UniProt ID **Q96VT3**), whose 3D structure is similar with a (β/α)_8_ barrel as the core (Supplementary Fig. S5). The alignment reveals chemically similar candidate active site residues and commonalities in ligand binding between the two glycoside hydrolase families, with a distinct **D**YG**E** consensus active site motif instead of the **E**WH**E** of GH99. *E. nidulans* GH71 does not have a residue analogous to Y189 of GH99 but a conserved and spatially close N74 may play a similar role in substrate binding.

### All non-bilaterian animal phyla except Ctenophora possess the endomannosidase

Analysis of my dataset showed that the all major animal clades, except Ctenophora, possess the GH99 endomannosidase protein. The sequences found in sponges, placozoans and cnidarians are always MANEA; indeed, I found no evidence of GH99-encoding gene duplications, or existence of MANEAL, in any major clades of non-gnathostomes. Sponges, placozoans and cnidarians usually possess one copy of the gene per species.

Strikingly, and contrary to previous claims^33,34^, endomannosidases were found in all four classes of sponges. While quite divergent, GH99 sequences from calcareous sponges (Calcarea) in most phylogenetic reconstructions form a sister group to sequences from the Homoscleromorpha sponge class, which is concordant with canonical sponge phylogeny^50^. In a number of demosponges and homoscleromorphs more than one copy per species was found. In *Stylissa carteri* and *Halichondria panicea* they are isoforms of the same gene but in the demosponges *Cymbastela concentrica* and *Spongia officinalis* analysis of a multiple sequence alignment (see Data Availability Statement) suggests they are products of different genes. In summary, while it also may have been lost in individual lineages, the endomannosidase activity seems to be a fixture among sponges, placozoans and cnidarians (except the highly specialized, parasitic myxozoans, see Supplementary File S6), but ctenophores do not possess it.

I detected one potential instance of lateral gene transfer from an ancestral Cnidarian/Placozoan to a common ancestor of two algae from the group Chattonellales: *Heterosigma akashiwo* and *Chattonella subsalsa* (Supplementary Fig. S1). Sequences from these species form a well-supported clade and the algal GH99 proteins contain the conserved animal residue Y189, while other sequences from ochrophytes mostly do not. This gene transfer could have occurred between a hypothetical ancestor of both Placozoa and Cnidaria^51^ and a common ancestor of Chattonellales. Further studies and timed phylogenies of Raphidophyceae are needed to investigate this possibility.

### All major clades of bilaterians except the sampled Xenacoelomorpha possess the endomannosidase

The average fraction of species that contained one or more 95% identity-clustered endomannosidase sequences in LukProt was similar in non-chordate bilaterian clades: 70-100% in arthropods, lophotrochozoans, ambulacrarians and non-vertebrate chordates, lower in nematodes (40%) and 0 in xenacoelomorphs. The number of Xenacoelomorpha taxa sampled was 3, too low to exclude the possibility of the gene existing in other xenacoelomorphs (Fig. 4).

**Fig. 4:**
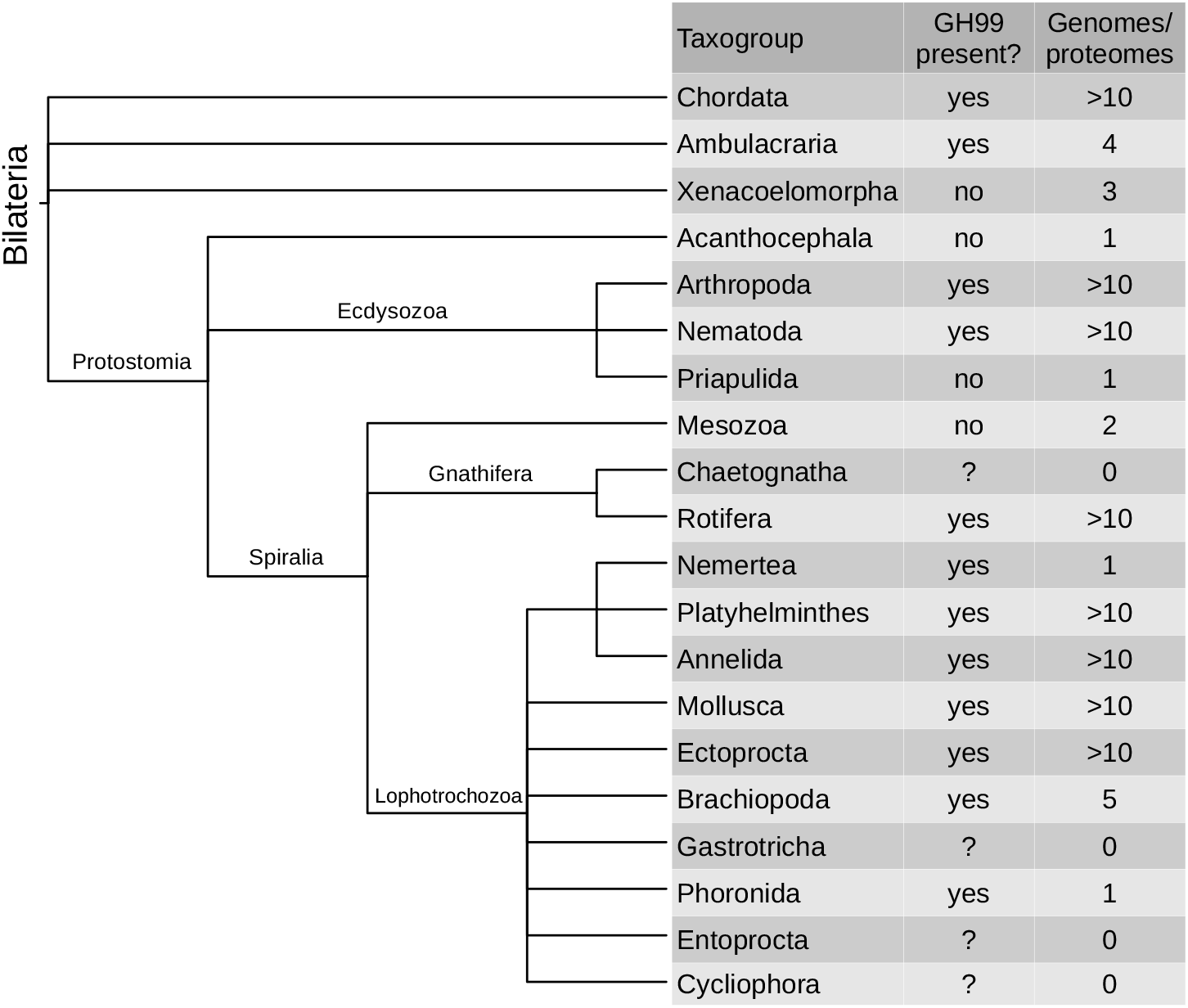
Cladogram and presence/absence analysis of the GH99 domain in bilaterian taxa, with emphasis on non-chordates. Clades are based on ^52^ and information from NCBI Taxonomy^53^.

By extending the search to all sequences available in the NCBI “nr” database, as well as TBLASTN searches on genomes for which predicted proteins are not available (Supplementary File S6) I surveyed a diverse sampling of protostomes. The single acanthocephalan genome did not contain sequences encoding for the GH99 domain. Rotifers have the domain but there are no genomes available of their probable sister clade^52^, Chaetognatha. Among lophotrochozoans, all clades with a genome available have GH99. The ecdysozoan clades that contain GH99 proteins are Nematoda (Enoplea and Chromadorea), Pancrustacea (crustaceans, ostracods, branchiopods and hexapods) and sequences from Diptera formed two distinct clades in the complete phylogeny (Supplementary Fig. S1). Both urochordates and hemichordates (Ambulacraria) possess the protein, as well as all the surveyed chordate clades. In summary, endomannosidase is widely distributed in bilaterians and only select clades do not contain any species with it.

### MANEA and MANEAL originated in the second round of vertebrate whole genome duplication

The timing and character of the two genome duplications that happened early in vertebrate evolution^54^ have been resolved recently via an analysis of deeply conserved synteny^55^. The 17 discovered LGs (linkage groups) were later revised to 18^56,51^ present in the ancestral chordate, but 17 were present in a proto-vertebrate immediately before the first whole genome duplication (1R). 1R happened in an ancestor of all extant verterbates (but not in nonvertebrate chordates: lancelets and tunicates). The second duplication (2R) took place in an ancestor of all jawed vertebrates (gnathostomes). While 1R (an autotetraploidy) led to symmetrical gene loss, 2R was a result of hybridization (allotetraploidy) of organisms α and β; the subsequent diploidization was asymmetrical. Taxonomic surveys of GH99 family proteins largely agree with this model: non-gnathostomes possess one copy of the EM protein, whereas most gnathostomes possess two copies: MANEA and MANEAL. (Supplementary Fig. S2, S3, coloring scheme: Supplementary Fig. S13)

A closer examination of the data published by Simakov et al.^55^ reveals that *MANEA* gene was located on the ancestral protovertebrate chromosome CLGK. *Branchiostoma floridae* chromosome BFL9 is the direct descendant of CLGK and in this amphioxus MANEA is localized there. The chromosomal locations of *MANEA* and *MANEAL* in other surveyed species (*Homo sapiens, Gallus gallus, Xenopus tropicalis*) help triangulate the origins of MANEA and MANEAL paralogs: MANEA comes from organism α and MANEAL from organism β. The α:β gene retention ratio is on average above 2^55^. (Supplementary File S7)

Members of the clade of cartilaginous fishes (Chondrichthyes), which diverged from other jawed vertebrates early, all have both MANEA and MANEAL. Among ray-finned fishes, non-teleosts have two proteins (MANEA and MANEAL) and teleosts from the Clupeocephala supercohort usually have three. The third GH99 protein from Clupeocephala consistently forms a separate clade, sister to the MANEAL clade (Supplementary Fig. S2). This strongly suggests that it is a paralog of MANEAL specific to clupeocephalan fishes. Taking this into account, I would like to name the clupeocephalan-specific paralog of *MANEAL* as *CMANEAL* (Clupeocephala *MANEAL*) and keep *MANEAL* as the name of the gene which retained more ancestral features. This pattern of paralogy is consistent with a previous studies which found that clupeocephalans have their rhodopsin^57^, and aromatase^58^ genes present in more copies in comparison to other teleosts, even though the teleost-specific genome duplication (3R) occurred in the common ancestor of Teleostei^59–62^. I detected no further duplications of endomannosidase retained in major vertebrate clades.

Phylogenetic analysis mentioned above unambiguously places sequences in the MANEA, MANEAL or CMANEAL clades. As automatic gene annotation is especially prone to errors resulting from gene paralogy, many genes that encode MANEA have been mislabeled as MANEAL. Supplementary File S8 contains a list of misclassified endomannosidase proteins and Supplementary File S9 contains a list of all classifications of vertebrate GH99 proteins. Supplementary File S12 contains lists of vertebrate species with only MANEA, only MANEAL/CMANEAL or with both.

### In vertebrates MANEA is rarely lost but MANEAL loss is common

Analysis of gene/protein presence in vertebrates showed that *MANEA* is hardly ever lost in this group. Uniquely, the only vertebrates that lost *MANEA* are the bowfin *Amia calva* and the albatross *Thalassarche chlororhynchos* in whose genomes this gene could not be detected. The catalytic activity of an endomannosidase seems to be essential for vertebrate survival, as both of these species have *MANEAL*.

*MANEAL*, in turn, seems to be more dispensable. In clupeocephalan fish, whose last common ancestor had CMANEAL, large clades acquired predicted inactivating mutations or gene losses. Cypriniformes, an order of Otomorpha, lost MANEAL but retained CMANEAL (Fig. 5A, Supplementary Fig. S2). In a major clade of clupeocephalans, Acanthomorpha, CMANEAL proteins suffered a probable loss of function mutation similar to GH99 inactive variants described earlier^21,40,63^ (E404Q). Their MANEAL proteins were consistently were active variants (E404), suggesting that MANEAL is catalytically active in these species but CMANEAL may not be.

**Fig. 5:**
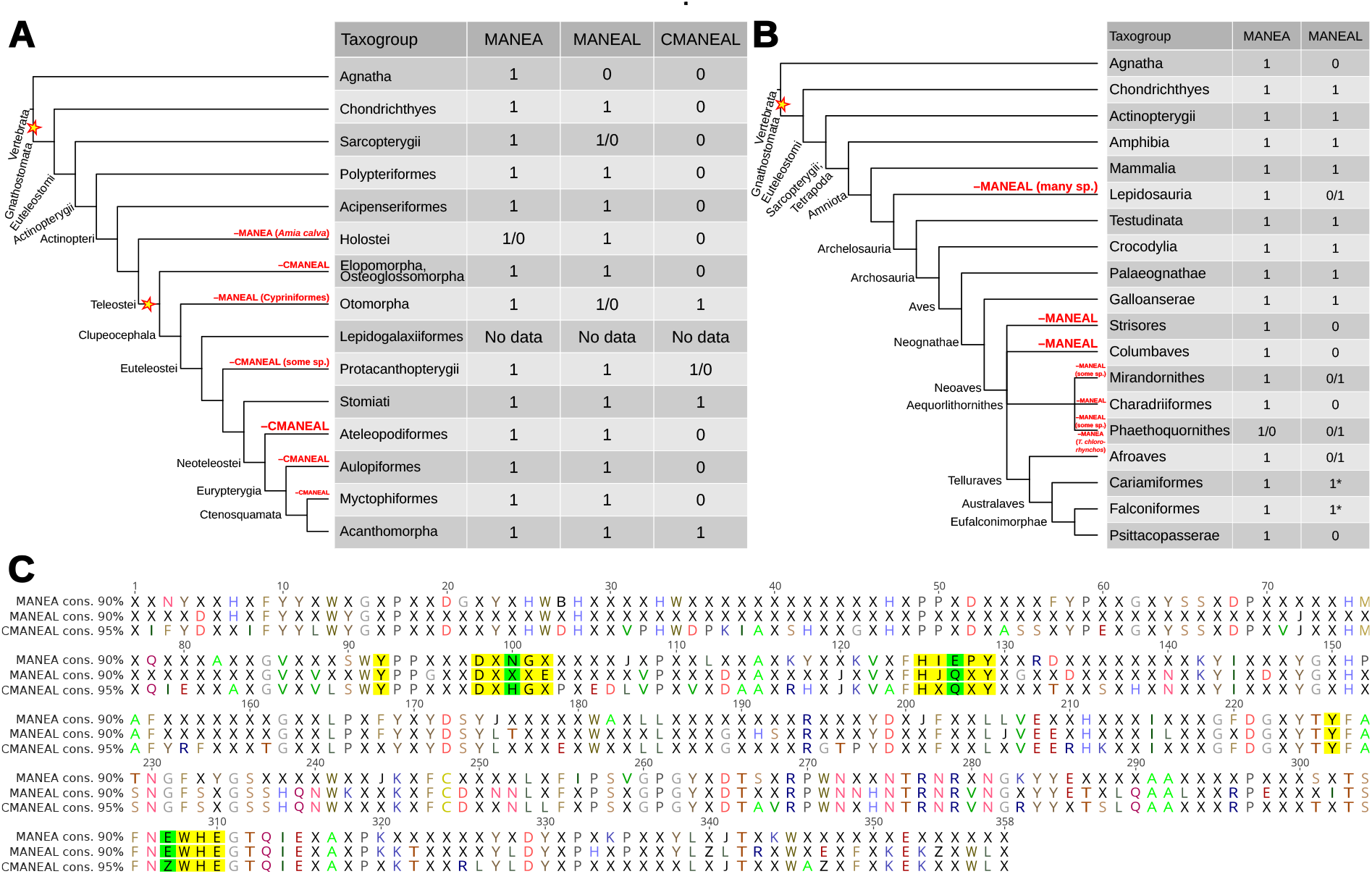
Analysis of vertebrate MANEA, MANEAL and CMANEAL: presence/absence and consensus (cons.) sequences. (A) Actinopterygii-centric cladogram, (B) Sarcopterygii-centric cladogram, (C) Alignment of catalytic domain consensus sequences (at 90% or 95% identity). Asterisks denote probable instances of pseudogenization and stars signify gene duplication events. The only 2 instances of MANEA loss in vertebrates are marked in red. In (C) the 5 investigated motifs are highlighted in yellow and crucial differences between them in the three orthologs are higlighted in green. Catalytic domain residue numbering is shifted by −97 with respect to human MANEA. For full phylogenies refer to Supplementary Fig. S2 and S3. Simplified fish phylogeny based on ^61^ and bird phylogeny based on ^64,65^.

TBLASTN searches of the three available genome assemblies of Acanthomorpha cousins within the clade Neoteleostei (Ateleopodiformes, Aulopiformes, Myctophiformes) suggested that only vestiges of CMANEAL are present in their genomes but they all seem to retain MANEA and MANEAL. These clades are likely to not produce an active form of CMANEAL protein. However, as the branches in the CMANEAL subtree are not long (Supplementary Fig. S2), it is possible that the protein gained a different function in these species – short branches suggest some selective pressure against random mutations, which would have been seen if the protein was dispensable and pseudogenized. Another consistent feature of CMANEAL is that the consensus sequence of the −2 motif is D[D/E] **H**GE instead of D[D/E]**N**GE present in other animal GH99 proteins (Fig. 5C). As residue N197 forms a hydrogen bond with the O4 of the −2 glucose^41^ (Fig. 3), the N197H variation might change the substrate binding affinity, possibly increasing it, as the electronegativity difference would be greater in case of the NH-O bond than in case of the OH-O bond. Interestingly, an N197H MANEAL protein variant is present in orders Charadiformes and Siluriformes, all within the Otomorpha clade that also harbors Cypriniformes in which MANEAL was lost (Supplementary Fig. S2). The inactivating E404Q mutation in CMANEAL did not occur in these two orders.

The loss of MANEAL also happened multiple times in various lineages of tetrapods. In Amphibia (amphibians) all major orders possess both genes. However, in squamates, the largest order of lizards, MANEAL appears to have been independently lost in Gekkota, Pleurodonta and Serpentes (snakes; with a caveat that there are no genomic data available for Leptotyphlopidae, Typhlopoidea and Amerophidia) (Supplementary Fig. S3). The overall availability of sequencing data in lizards is low and these results are tentative. The only living non-squamate reptile, the tuatara, possesses both MANEA and MANEAL.

Among birds, MANEAL losses are common but their pattern is more scattered. The major bird clades lacking MANEAL are Strisores and Columbaves. All species within Galloanserae contain both proteins. Among Neoaves, within the large waterbird clade Aequorlithornithes, both proteins are retained in some species in the subclades Mirandornithes and Phaethoquornithes, but the major clade Charadriiformes lacks MANEAL (Fig. 5B). In Telluraves (a Neoaves subclade), MANEAL was lost in Psittacopasserae (passerines + parrots), but in their closest cousins, falcons and cariamiformes (*Cariama cristata* and *Chunga burmeisteri*) a MANEAL sequences with mutations in the active site were detected.

For example, falcon MANEAL predicted proteins are N-terminally truncated and have a mutated active site (**E**WH**E**→**K**WH**E**). MANEAL therefore probably pseudogenized is the last common ancestor of Australaves and was subsequently lost from the genome in the last common ancestor of Psittacopasserae. Among Afroaves many species lost MANEAL, but some (*Indicator maculatus, Ramphastos sulfuratus*) retained it.

Tetrapod phylogenies also uncovered that in the bird and lizard species which retained MANEAL, the protein underwent faster change than MANEA. While the MANEA branch is largely representative of true phylogenetic relationships between species (Supplementary Fig. S3), the MANEAL clade is not: squamate MANEAL proteins were incorrectly placed inside the Archelosauria clade with a high UFBoot2 support of 97. The crocodile MANEAL clade was artefactually placed inside the bird MANEAL clade (UFBoot2: 62) and the branches were long. The probable falcon pseudogenic MANEAL proteins formed a tight cluster on a particularly long branch. Many other archelosaurian MANEAL protein sequences are found on long branches, supporting the pseudogenization hypothesis (Supplementary Fig. S3).

My analysis of the CMANEAL clade (Supplementary Fig. S2) suggests that despite suffering a probable inactivating E404Q mutation, in most species the gene likely did not pseudogenize. Such a process would have been visible as an abundance of long branches resulting from sequences that acquired random mutations. In Fig. 5 I present an alignment of the consensus catalytic domains of animal vertebrate MANEA, MANEAL and CMANEAL. For each ohnolog, I prepared a separate catalytic domain profile, found the hits using HMMER and plotted the difference in e-value for each hit that all models detected. I obtained easily identifiable clusters representing MANEA (together with ancestral sequences) MANEAL and CMANEAL (Supplementary Fig. S3).

Further, my results showed that all mammals have both MANEA and MANEAL. In the cases where both of the genes were labeled with the same name (MANEAL), through phylogenetic analysis I found these to all be annotation errors. A number of mammalian genome annotations (12 species) suggested they only possess MANEA and 6 other annotations state MANEAL only. I conducted TBLASTN searches of each of these assemblies (Supplementary File S6) and BLASTP searches of the translated protein models to test this. In every case I found both MANEA and MANEAL.

## Discussion

In this work I described the evolution of motifs in GH99 domain proteins, its abundance in various taxonomic groups placed in their phylogenetic context, and uncovered a novel *MANEAL* paralog in animals. I tested the conclusions of both of the initial attempts at the analysis of endomannosidase origins: a 1997 phylogenetic survey by Dairaku and Spiro^33^ and Sobala 2018^34^ and found them incomplete and incorrect. The authors of the early taxonomic survey did not enjoy the current abundance of sequencing data and sampled only 20 taxa, while in the current analysis I included hundreds of species. While some protostome GH99 proteins are often quite divergent from vertebrate endomannosidases and might have evolved different activities, in this study I focused on prevalence of the GH99 domain only and can only infer activity conservation. To directly investigate enzymatic activity, biochemical studies comparing sequences with different active site motifs are needed. The specific claims refuted by this analysis are: (1) that endomannosidase is a recent addition to the eukaryotic N-glycan processing machinery, (2) that it is limited in distribution to chordates with the exception of molluscs. Contrary to Sobala^34^, it is also evident that MANEA and MANEAL genes are present in all jawed vertebrates – not only in bony vertebrates.

Currently, public databases are plagued by extensive errors in paralogous gene annotation, which propagate to other species, lowering the collective quality of publicly accessible sequence data. The analyses presented here should result in corrections to databases and a recognition of a new family of vertebrate MANEAL sequences – CMANEAL – from teleost fishes in which the gene duplicated (supercohort Clupeocephala). While this study is not broad enough to test this, previous evidence^57,58^ suggests that CMANEAL was lost in Elopomorpha and Osteoglossomorpha, rather than duplicated in Clupeocephala. I predict that the two almost universal substitutions that characterize CMANEAL, **E**404**Q** and **N**197**H,** caused loss of its activity and an increase in its binding affinity. This suggests that the protein evolved a separate function, possibly becoming a lectin. Only kinetics studies can confirm or refute this.

It is still unclear why the GlcMan3-specific Y189 form of the endomannosidase is conserved in almost all animals, while throughout eukaryotes the prevailing form is W189. The precursor glycan, while shorter in some species, has the Glc3Man9GlcNAc2 structure in groups as distant as plants and animals^29,30^. Its truncation is though to be an effect of secondary losses of glycosyltransferases within eukaryotes^30^. An attractive proposition is that within filozoans (see Fig. 2), the increasing complexity of life histories associated with predatory lifestyles^66–68^ required higher quality control mechanisms. These, in turn, potentiated filozoans and choanoflagellates to subsequently evolve clonal lifestyles and multicellularity. In multicellular organisms cells must communicate their identity to others precisely, otherwise they could be recognized as non-self and expelled or killed. N-glycans on membrane-anchored proteins may form such signaling tags. In addition, the N-glycosylation machinery is used for protein folding quality control and some misfolded proteins could also be recognized as non-self. In this view, the evolution of multicellularity depended on and potentiated the selection of well-defined (identity-signaling) cell surfaces, which in turn depended on effective cellular mechanisms of quality control. While such mechanisms are expensive for single cells in terms of energy, they nevertheless have a potential to confer selective advantages to assemblies of cells (organisms) that possesses these traits. Hypothetically, the GH99 endomannosidase could be an important factor in such processes as it enables further processing of more N-glycans in the Golgi. The case of ctenophores not having the endomannosidase at all speaks to their uniqueness in the animal kingdom. It is likely that they evolved a different mechanism of dealing with precise glycan editing.

I uncover a large expansion of GH99 family proteins in diatoms (Supplementary Fig. S1). The only diatom whose N-glycome was studied to date, *Phaeodactylum tricornutum*^69,70^, happens to have only one GH99 protein (clustering within Diatomeae GH99 major clade 2) which was omitted from the bioinformatic analyses in these studies. Such an expansion implies functional diversification of GH99 proteins in these species.

Building on the understanding of GH99 structure and evolution, I report putative ligand binding residues and active site residues of a protein from a family related to GH99, a fungal GH71 mutanase. The active site residues of these proteins were unknown to date. Distant but detectable evolutionary relationships such as these can be used to study glycoside hydrolases and generate hypotheses about their mechanism of action.

A limitation of the current study is that pseudogenization is investigated only indirectly (through phylogeny analysis and sequence comparison). Due to the presence of transcriptomes in the LukProt database, pseudogenes might be visible by being expressed at a low level and then assembled and transcribed *in silico* into partial proteins. This would increase the apparent number of species containing both MANEA and MANEAL. Another limitation is that in cases where TBLASTN searches of genome assemblies were ran to confirm protein BLAST results, any indication of MANEA or MANEAL in the genome was counted as their presence. This could have inflated the number of species and clades in which the sequences were detected. Such an approach was selected in order to understand broad patterns of endomannosidase evolution and not its evolution in individual taxa. I also did not directly investigate the functions of CMANEAL.

Another potential weakness of the work is the high number of sequences used to produce single-gene phylogenies, which in some cases led to overparametrization. The affected single gene phylogenies were treated as guides only, with clades concordant with species phylogeny being treated as more informative than those which grouped together unrelated species.

As bacteria, archaea and most clades of eukaryotes possess GH99 domain proteins, the last eukaryotic common ancestor probably possessed a gene encoding for a protein with this domain. The form of the enzyme containing all the 5 sequence motifs typical to animal endomannosidases emerged early, probably in a common ancestor of all eukaryotes. It would be a remarkable case of convergent evolution if they appeared separately in most Diaphoretickes and Podiata. Specificity towards GluMan3 (Y189) became fixed in the ancestral choanozoan, and is conserved in most species from this clade. The last common ancestor of all animals posessed one copy of the gene encoding MANEA and all major animal clades, except the ctenophores, inherited it. The endomannosidase activity is essential for every vertebrate and for many invertebrates. Patterns of endomannosidase-like gene gains and losses suggest that its activity is essential for most animals, but the ohnolog of *MANEA* – *MANEAL* – and the ohnolog of the latter, here named *CMANEAL*, are often lost or acquire inactivating mutations. *CMANEAL* was gained only by clupeocephalan fishes after the cladespecific gene duplication. It is not known whether variants of this full-length GH99 domain protein with the E404Q mutation have enzymatic activity: they might have evolved into a lectin.

In summary, the findings presented here allowed to track the evolutionary story of GH99 domain usage by various groups of organisms and speculate on the possible importance of endomannosidase activity in animal multicellularity. I hope that this study will clear the picture of the endomannosidase protein family evolution and provide the reader with an understanding of the functional importance of this enzymatic activity. Finally, misconceptions stemming from animal-centric and model organism-centric approaches, are difficult to dispel and lead to erroneous assumptions which may be reproduced in literature for decades. I would like to encourage the reader to always consider the full tree of life when looking for origins of specific cellular activities^71^.

## Materials and Methods

The LukProt database version 1.4.1^72^ was used in all local BLAST+^73^ searches (blastp, tblastn). Remote blastp searches were done using NCBI BLAST online service^74^ against the nr database and remote tblastn searches were performed using “BLAST Genomes” option. Bacterial, archaeal and metazoan GH99 sequences from the nr database were retrieved using blastp against the nr database using UniProt ID **Q5SRI9** (human MANEA) with max_target_seqs 5000 and an e-value of 1 × 10^-4^. Additionally, picozoan GH99 sequences were found using EukProt v3^37^ BLAST server (https://evocellbio.com/SAGdb/EukProt/) by running blastp of Q5SRI9 as query against all Picozoa. Unpublished sequences from a filasterean *Txikispora philomaios*^75^ were searched using the same method. The user interface for local BLAST+ (v2.13.0) was SequenceServer^76^. The software used for sequence alignment was MAFFT v7.508^77^ (L-INS-I mode), MUSCLE v5.1^78^ or Famsa v2.2.2^79^. Alignments were then trimmed using trimAl^80^ (gap threshold 0.05 to 0.1) and, in cases explained in the associated Zenodo dataset (doi:10.5281/zenodo.7470560), manually in Geneious Prime v.2022.2.2 (https://www.geneious.com). To search LukProt with a GH99 domain hidden markov model (Pfam accession **PF16317.7**) and with GH99 profiles generated in this work, HMMER v3.3.2 (http://hmmer.org) software was used. Phylogenies were prepared using IQ-TREE 2.2.0-beta^81–83^ and visualized in Figtree v1.4.4 (http://tree.bio.ed.ac.uk/software/figtree/) or Dendroscope v3.8.4^84^. Unless otherwise noted, the default options changed in IQ-TREE were: --msub nuclear -alrt 5000 -bb 5000 -abayes -nm 10000. Sequences were clustered at various identities (50-100%) using CD-HIT^85,86^ (command example for clustering at 95% identity: cd-hit -s 0.5 -g 1 -d 0 -T 6 -M 16000 -c 0.95). Signal peptide searches were carried out using with SignalP 6.0^87^. In some cases I found very divergent GH99 proteins, which may or may not be artifacts of *in silico* transcript reconstruction and translation. Sequences that were obviously erroneously translated were excluded from single gene phylogenies. The list of excluded sequences with the reasons can be found in Supplementary File S10. Other sequence handling tasks were accomplished using SeqKit^88^, GNU coreutils and Geneious Prime. For details of individual alignments and phylogeny reconstructions the reader is directed to the README files in the aforementioned Zenodo dataset.

## Supporting information

Supplementary Data Descriptions

## Funding

This work was supported by National Science Centre of Poland as part of the grant SONATINA no. 2020/36/C/NZ8/00081 entitled “The role of glycosylation in the emergence of animal multicellularity”.

## Acknowledgements

I would like to acknowledge Marcin Czerwiński and Iñaki Ruiz-Trillo for a critical reading of the manuscript. I thank Michelle Leger and Koryu Kin for advice with bioinformatics and indispensable comments which improved the manuscript. Part of the calculations for this project were done using the Marvin cluster of Universitat Pompeu Fabra, Barcelona, Spain.

## Abbreviations

1R/2R/3R: first/second/third round of vertebrate genome duplications
CAZy: Carbohydrate Active Enzyme database
CLGK: conserved linkage group K
*CMANEAL*: Clupeocephala *MANEAL*
cons.: consensus
ECM: extracellular matrix
Fig.: Figure
Gal: galactose
GH: glycoside hydrolase
Glc: glucose
GlcNAc: *N*-acetylglucosamine
HMM: hidden Markov model
Man: mannose
NCBI: National Center for Biotechnology Information
OST: oligosaccharyltransferase
sp.: species

## Data Availability Statement

Source data, scripts and supporting data can be found in the associated Zenodo repository: https://doi.org/10.5281/zenodo.7470560.

## Notes

https://doi.org/10.5281/zenodo.7470560

### Competing Interest Statement

The authors have declared no competing interest.

