## Supplementary Data Descriptions for "Evolution and phylogenetic distribution of *endo*-α-mannosidase"

Supplementary Fig. S1: Annotated phylogeny all eukaryotic GH99 sequences, decontaminated, clustered at 95% identity, prepared using iqtree2. Gnathostome clades are colored according to protein identity: blue – MANEA, red – MANEAL, orange – CMANEAL.

Supplementary Fig. S2: Annotated phylogeny of vertebrate sequences from LukProt and vertebrate but not tetrapod GH99 sequences found using BLAST of the NCBI nr database. Colors as in Supplementary Fig. S1.

Supplementary Fig. S3: Annotated phylogeny of vertebrate sequences from LukProt and tetrapod but not mammal GH99 sequences found using BLAST of the NCBI nr database. Colors as in Supplementary Fig. S1.

Supplementary Fig. S4: Evaluation of the specificity of various HMM profiles of endomannosidase protein. Each point represents a single sequence found by searching LukProt using all models and its placement is based on the log(evalue_model 1_/evalue_model 2_), such that the lower the value, the more likely the sequence is to be found by model 1. 0 represents equal probability. (A) MANEAL/CMANEAL versus MANEA/MANEAL. (B) MANEAL/CMANEAL versus opisthokont/animal. (C) MANEA/MANEAL versus animal/MANEA. (D) animal/MANEA versus opisthokont/animal. GH99 sequences from known genes were colored according to the legend.

Supplementary Fig. S5: 3D alignment of human MANEA structure (ice blue, PDB ID: **6ZFA**) and the AlphaFold model of *E. nidulans* GH71 mutanase (gold, UniProt ID: **Q96VT3**).

Supplementary File. S6: Data on further investigation of taxa not covered by the LukProt database but essential for finding gains and losses of the endomannosidase by various clades, categorized by taxogroup. These investigations were done mostly using genomic TBLASTN.

Supplementary File. S7: Loci of endomannosidase genes in various vertebrates and the resulting protovertebrate CLGK locus.

Supplementary File. S8: List of MANEA sequences misclassified as MANEAL in public databases.

Supplementary File. S9: Table of sequence identifiers of GH99 sequences, corresponding species and taxogroups and protein classification (MANEA/MANEAL/CMANEAL), as well as sequence cleaning status (retained, divergent or contamination).

Supplementary File. S10: A separate table summarizing phylogeny-based sequence cleaning.

Supplementary File. S11: Abundance of sequences and particular sequence features in GH99 proteins from various eukaryotic taxogroups. Abbreviations are explained in the file.

Supplementary File. S12: Lists of species from various taxogroups categorized by which proteins they harbor. Species highlighted blue were found in public databases to only have MANEA, species highlighted red were thought to only have MANEAL and non-highlighted species were though to contain both proteins. Species were moved after classification performed in this work but their highlighting was not changed to show current database errors.

Supplementary Fig. S13: Phylogeny coloring scheme used in all trees. The following clades have multiple colors in their subclades (listed from left to right): Nucletmycea^a^: Rotosphaerida, Fungi; Choanoflagellata^b^: Craspedida, Acanthoecida; Porifera^c^: Demospongiae, Hexactinellida, Homoscleromorpha, Calcarea; Cnidaria^d^: Cubozoa, Hydrozoa, Scyphozoa, Hexacorallia, Octocorallia, Myxozoa; Protostomia^e^: Nematoda, Arthropoda, Platyhelminthes, Mollusca, Annelida, other Lophotrochozoa.
