## Supplementary figures and images for "Evolution and phylogenetic distribution of *endo*-α-mannosidase"

### Supplementary Figure 1

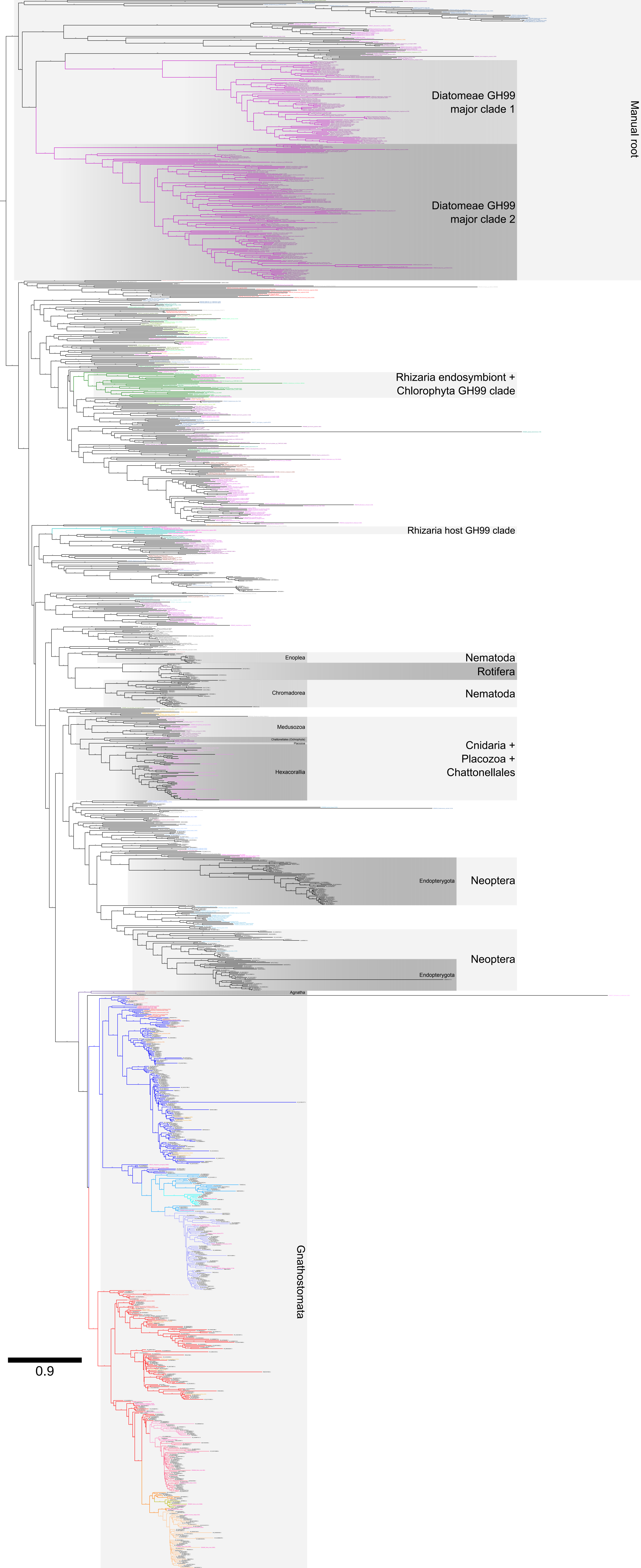

### Supplementary Figure 2

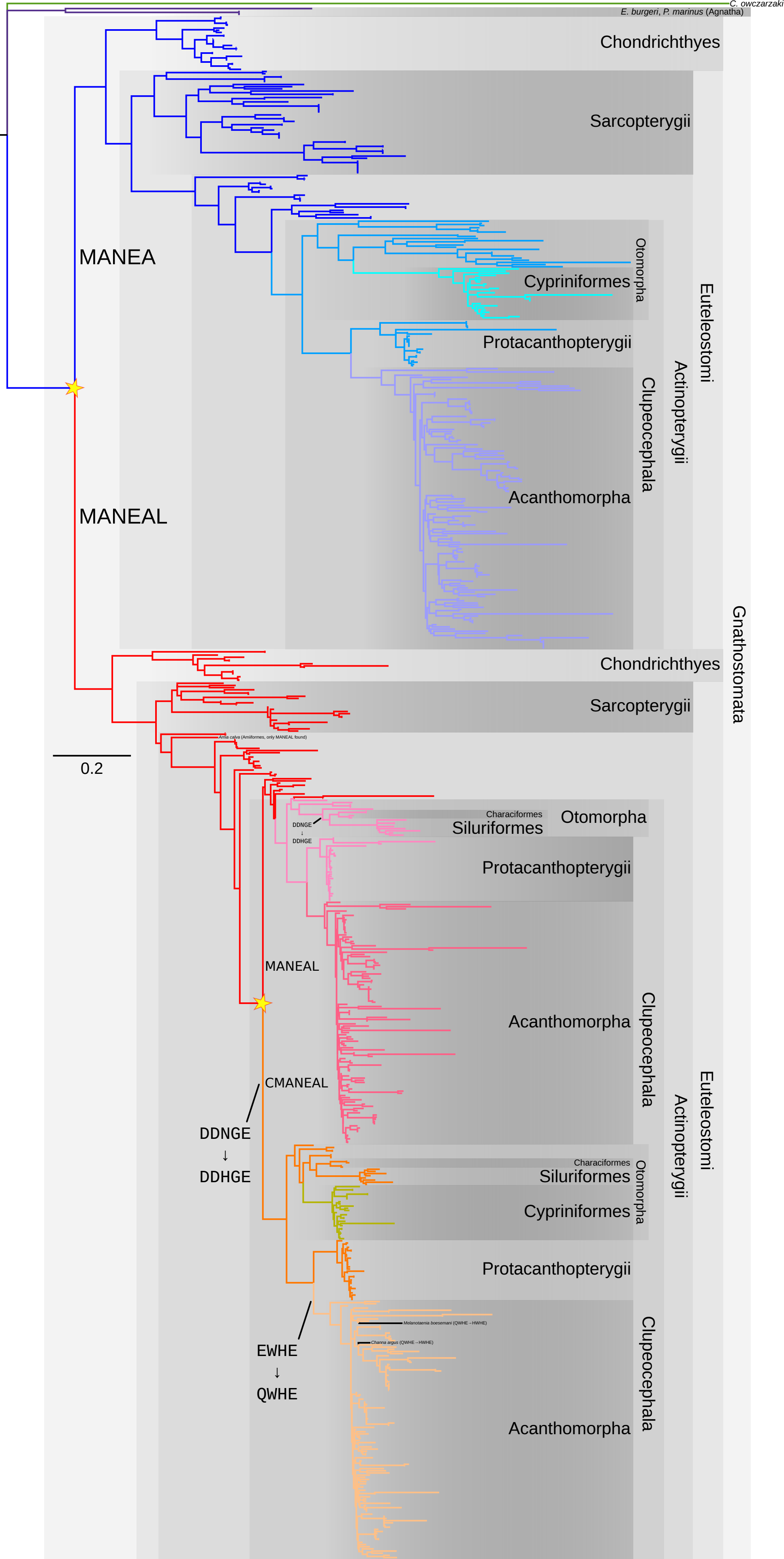

### Supplementary Figure 3

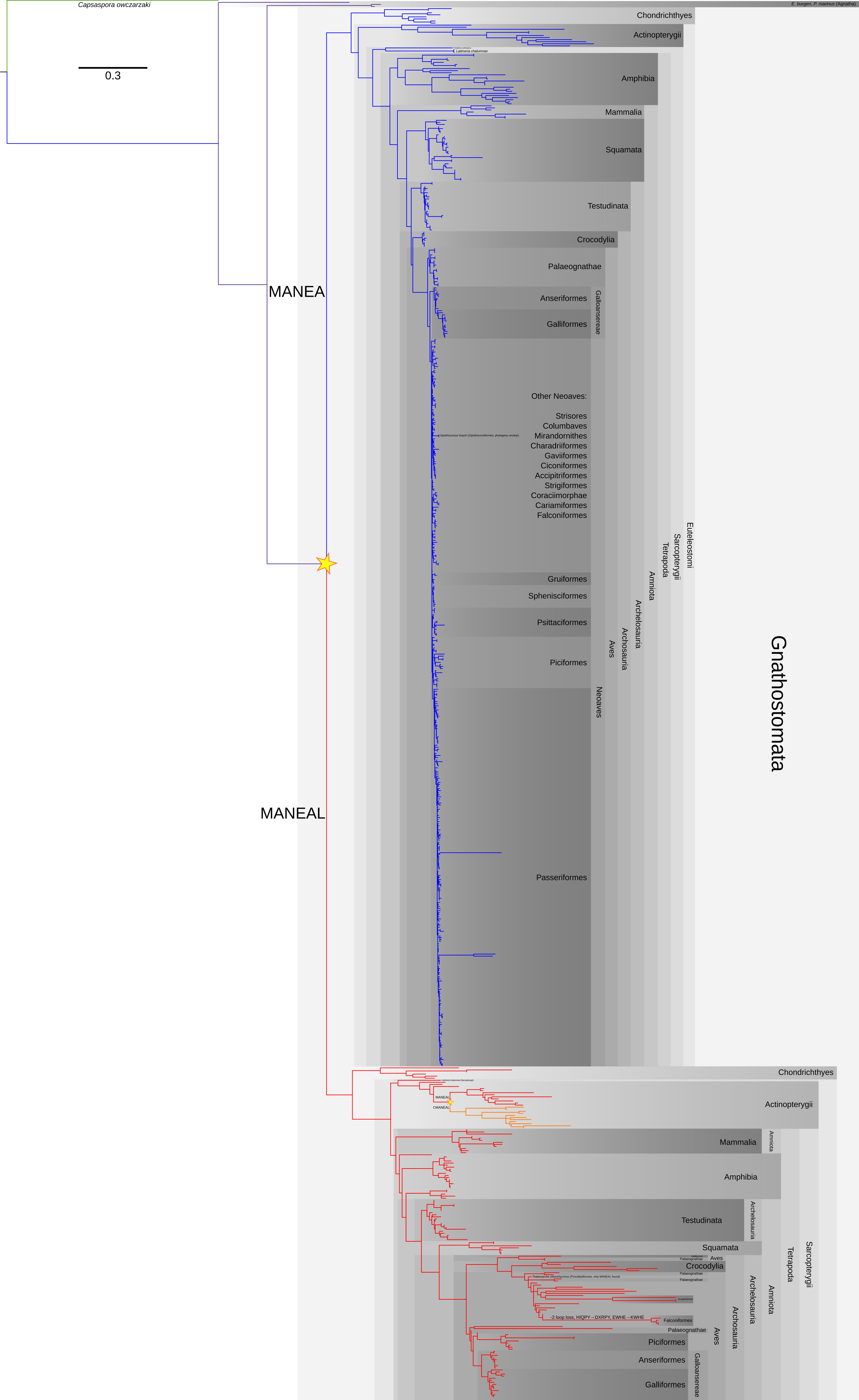

### Supplementary Figure 4

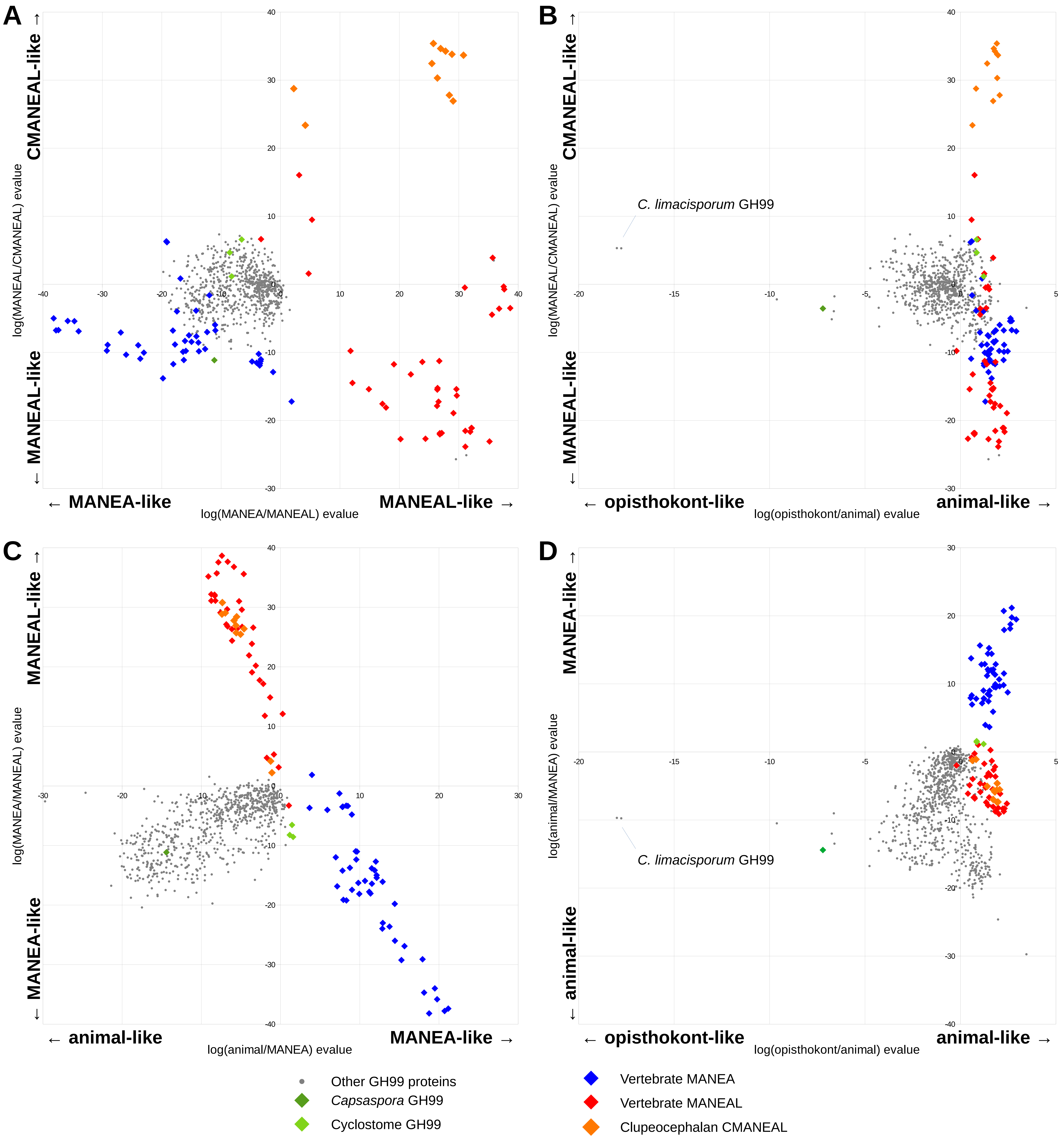

### Supplementary Figure 5

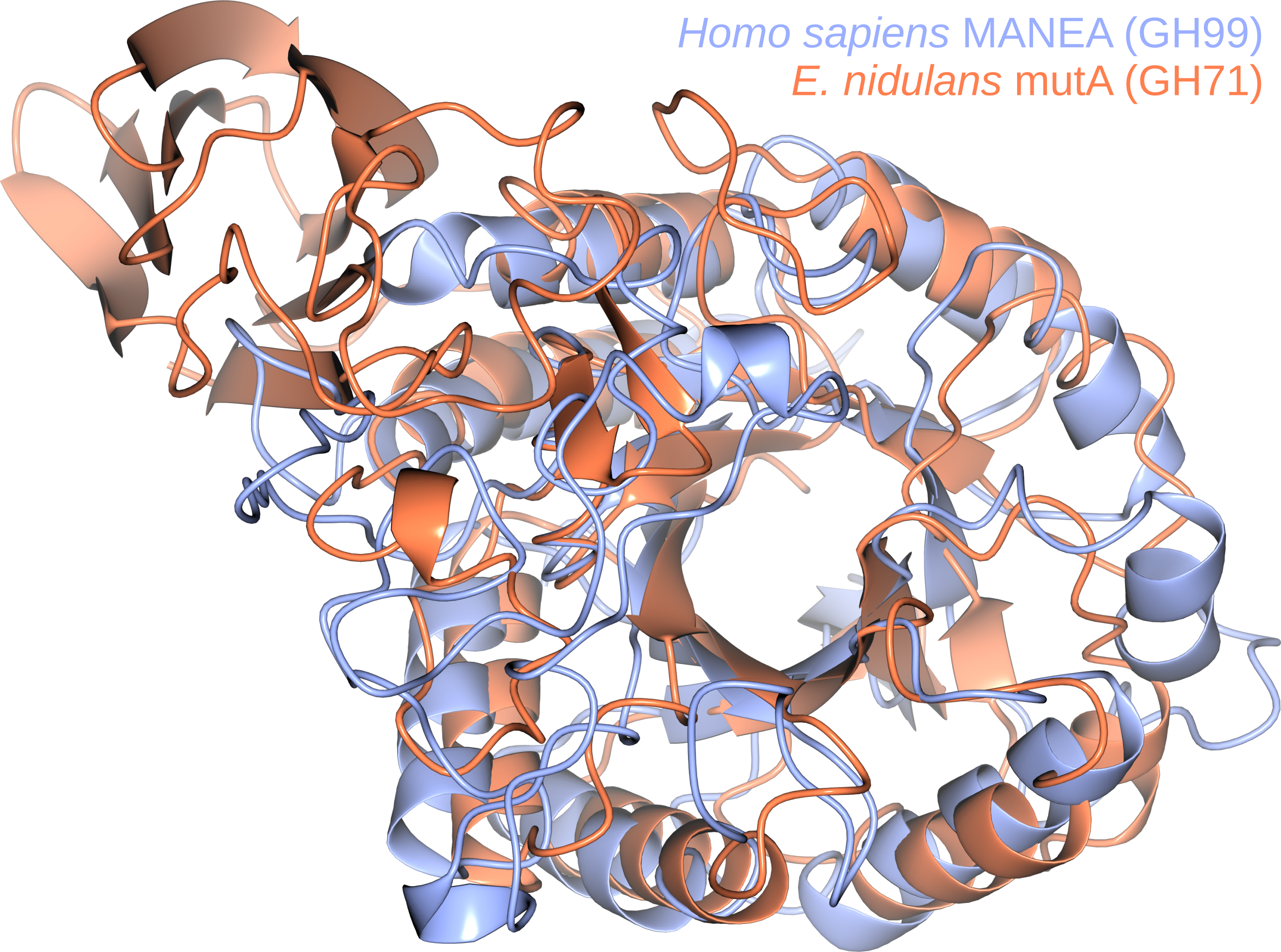

### Supplementary Figure 13

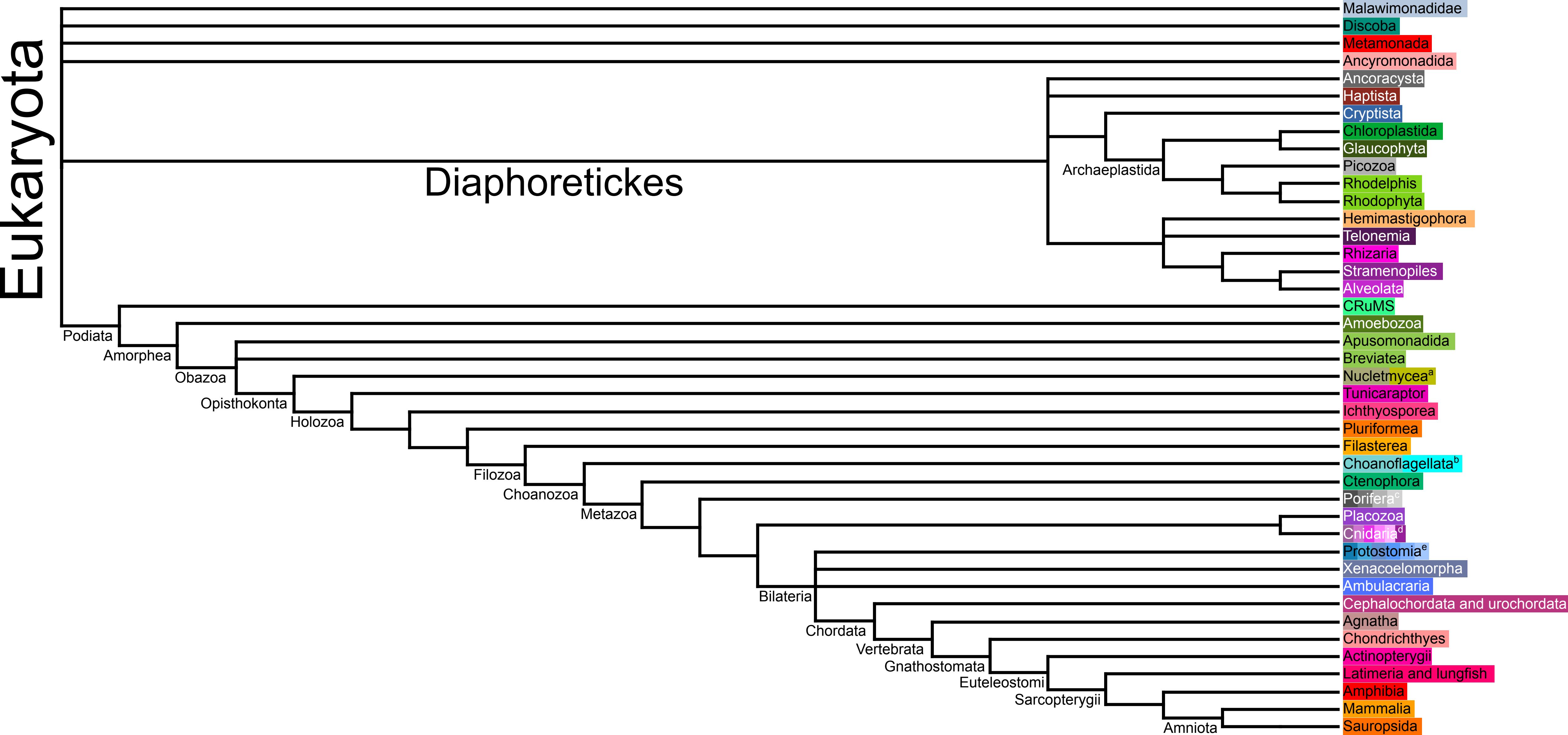
